# High-resolution cryoEM of nucleosomes in nuclear extracts of mammalian cells

**DOI:** 10.64898/2026.06.15.732463

**Authors:** De-Sheng Ker, Hedaya Aboalnaga, Luca Pellegrini

## Abstract

Frontier Structural Biology methods are transitioning from analysis of reconstituted macromolecular complexes *in vitro* to imaging of macromolecular assemblies within the physiological confines of the cell. Preparation of samples for *in situ* cryoEM analysis requires FIB milling or ultramicrotome sectioning, laborious and technically challenging procedures that are low-throughput and require a high degree of technical skills. We have devised a simple approach for cryoEM of nuclear macromolecular complexes that preserves to a high degree their physiological environment while removing the need for thin sectioning of the sample. The method requires only the preparation of nuclear extracts without additional purification or enrichment steps. We applied the method to obtain a 2.3 Å cryoEM structure of nucleosomes visualised directly in the nuclear lysate of human cells.

**Graphical abstract:** 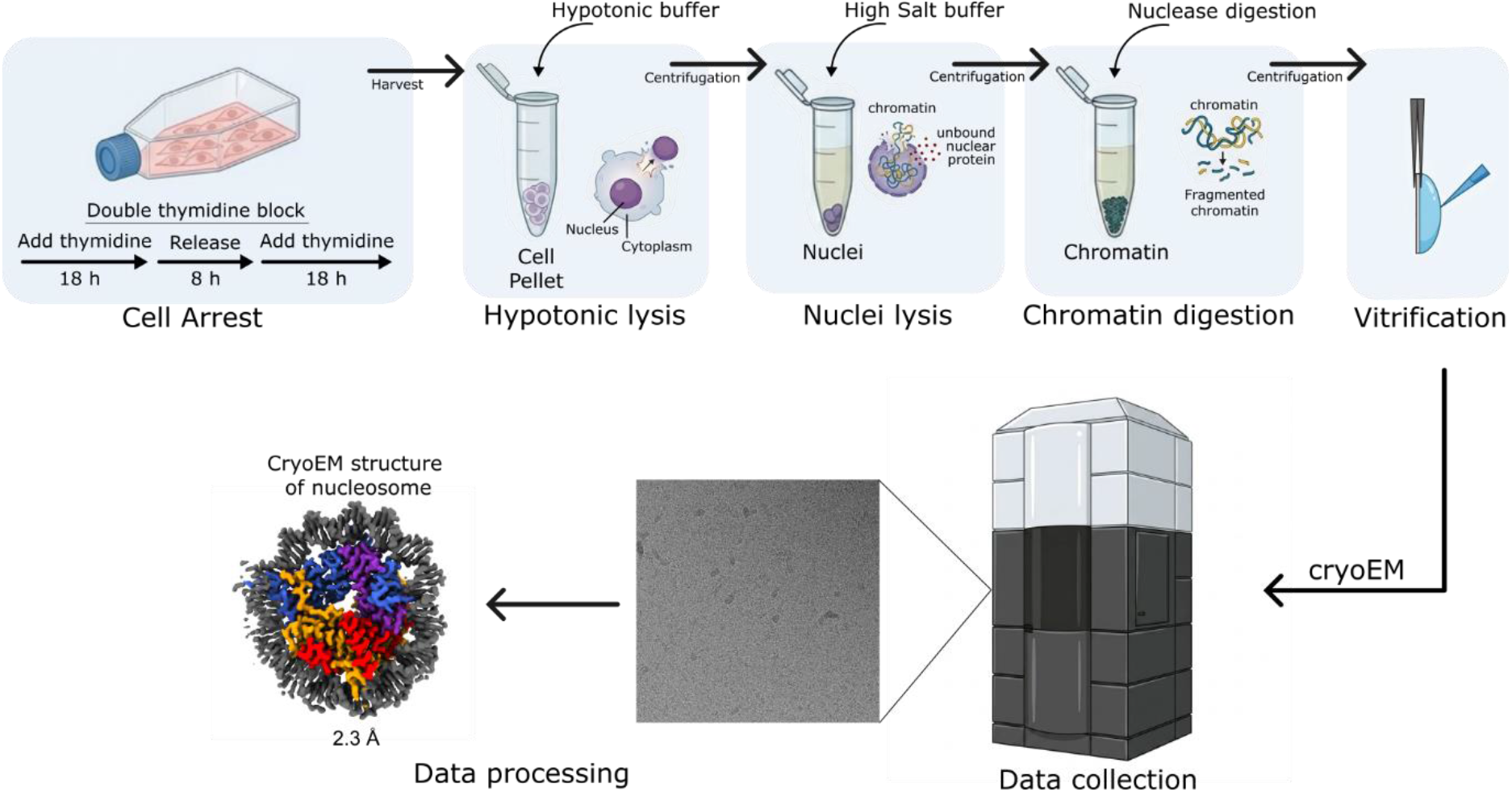

## Introduction

Recent technological advancements in Structural Biology have enabled the visualisation of ever larger macromolecular assemblies at near atomic resolution. The resolution revolution in cryo-electron microscopy (cryoEM) has combined improvements in hardware, such as the advent of direct-electron detectors, and image-processing software^1–3^ to help reveal mechanistic insights into fundamental processes such as RNA transcription^4^, ribosome biogenesis^5^, spliceosome assembly^6^, and flagellar motility^7^. However, achieving three-dimensional reconstruction at high resolution normally necessitates the biochemical reconstitution from purified components or isolation from the cellular environment, resulting in structural information that is void of its native context.

The frontier of the field has therefore gradually shifted toward *in situ* Structural Biology, in which macromolecular complexes are imaged directly within the native context of the cell. Cryo-electron tomography (cryoET), together with subtomogram averaging, has begun to yield macromolecular complexes in their native environments, capturing details such as the interactions with the membrane and cellular ultrastructure^8,9^. However, the transition from biochemical reconstitution to intracellular imaging introduces formidable technical barriers that need to be overcome. The principal obstacle is that most bacterial and all eukaryotic cells exceed the 300 nm thickness limit required for high resolution cryoEM imaging, thereby requiring sample thinning procedures such as focused ion beam (FIB) milling or cryo-ultramicrotomy^10,11^. Both methods are labour intensive, low throughput and demand specialized instrumentation and operator expertise. Consequently, *in situ* cryoEM remains currently restricted to a limited number of expert laboratories, restricting its broader adoption.

These constraints highlight the need for methodologies that, whilst preserving near physiological conditions of the sample, bypass the requirement for cell thinning procedures. The *ex vivo* application of cellular extracts directly onto the cryoEM grids offers an attractive alternative, as a compromise between the preparation of lamellae by sample thinning and a fully reconstituted biochemical system. Several groups have already demonstrated the feasibility of visualising macromolecules directly from cell lysates by cryoEM and, when combined with single particle analysis (SPA), achieving resolutions comparable to those obtained working with purified macromolecule complexes^12–14^.

The protein assemblies that maintain, duplicate and read the genetic information encoded in our DNA reside within the nucleus of eukaryotic cells. It is therefore important that we develop methods for imaging these nuclear assemblies in conditions that approximate as closely as possible their native state. Here we therefore extend the *ex vivo* approach to cryoEM imaging of nuclear extracts. We have devised a protocol that relies on a standard nuclear extraction preparation followed by plunge freezing for cryoEM analysis. The protocol does not require additional purification steps beyond the nuclear extraction, and the process from lysis to vitrification can be completed in less than two hours using standard cryoEM equipment.

We demonstrate the validity of this method by determining the cryoEM SPA structure at 2.3 Å resolution of the nucleosome core particle (NCP) directly from nuclear extracts of mammalian HEK293T cells. Our method provides a foundation for development of nuclear extraction protocols aimed at visualising macromolecular assemblies of interest directly within nuclear extracts.

## Materials and Methods

### Cell Culture and synchronization

HEK293T cells were grown in DMEM medium supplemented with 10% (v/v) fetal bovine serum (FBS) at 37 °C in a 5 % CO_2_ atmosphere. Cells were grown to 50 % confluence in T-25 flasks before synchronization. To synchronize the cells at the G1/S boundary, the cells were treated with 2.5 mM thymidine for 18 h, washed thrice with PBS and released for 8 h, followed by an additional thymidine block of 18 h. After synchronisation, cells were trypsinized and harvested by centrifugation at 500 x *g* for 5 min, and the cell pellets were washed once with PBS.

### Nuclei isolation

The cell pellet was lysed on ice for 10 min in 50 volumes of hypotonic buffer (10 mM HEPES pH 7.5, 10 mM NaCl, 0.1% (v/v) NP-40, 3 mM MgCl_2_) supplemented with 1 mM TCEP, 1x Phosphatase inhibitor Cocktail (Sigma-Aldrich, P5726) and 1x Protease inhibitor Cocktail (Sigma-Aldrich, P8340) solutions. Nuclei were pelleted by centrifugation at 800 x g for 5 min at 4 °C. The supernatant was removed, and the nuclei pellet was resuspended once with hypotonic buffer and divided into aliquots of 0.5 to 1 million cells for testing of lysis conditions.

### Nuclear extract preparation

Nuclear lysis optimization was achieved by means of detergent or high-salt conditions as described below (Figure 1). All buffers were supplemented with 1 mM TCEP, 1 mM ATP, 1x Phosphatase inhibitor Cocktail (Sigma-Aldrich, P5726) and 1x Protease inhibitor Cocktail (Sigma-Aldrich, P8340) solutions, immediately before use.

**Figure 1:**
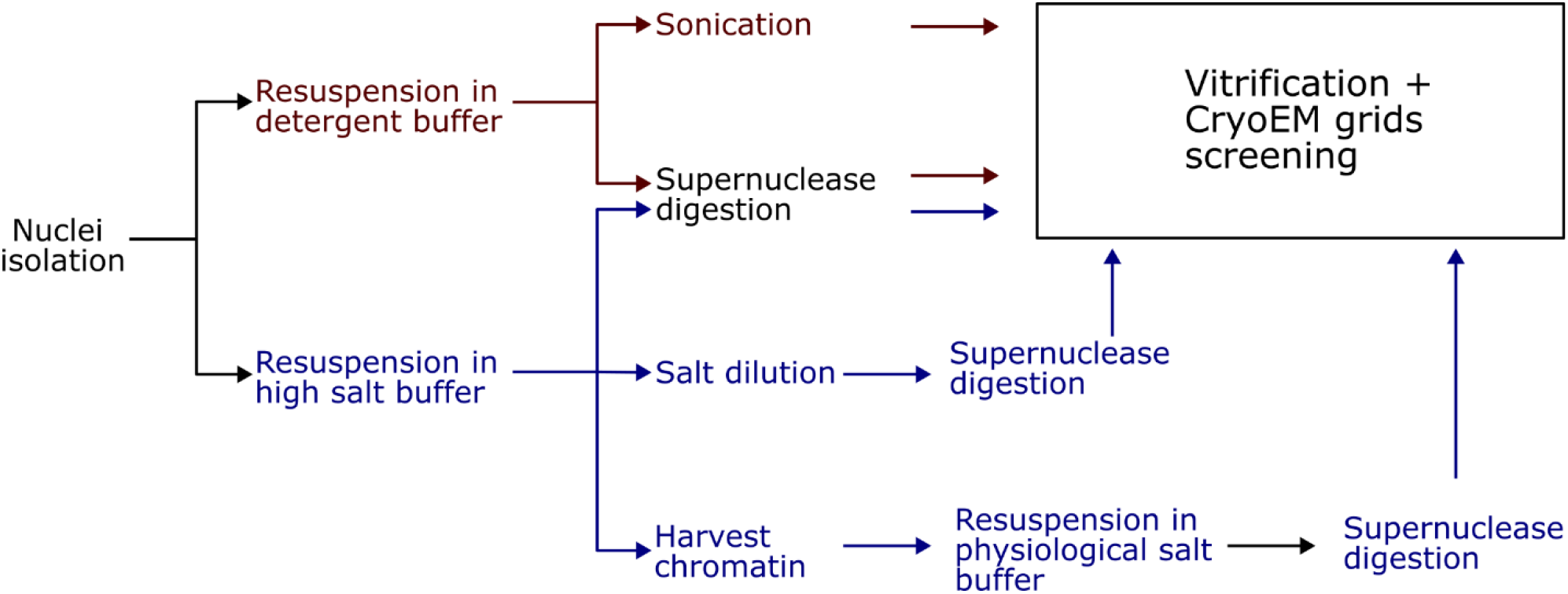
Nuclear lysis optimisation workflow for cryoEM sample preparation. Nuclei were isolated after hypotonic lysis and subjected to either detergent-based or high-salt-based lysis approaches. Chromatin was then fragmented using either sonication or SuperNuclease digestion prior vitrification on cryoEM grids for Talos screening.

#### Detergent-based nuclear lysis

The nuclei pellet was resuspended in 100 μL of buffer: 20 mM HEPES pH 7.5, 50 mM NaCl, 0.1% (v/v) Triton X. The chromatin was fragmented using either an enzymatic or a mechanical approach. For enzymatic fragmentation, the nuclei were incubated with 1 unit of SuperNuclease (SinoBiological, Cat: SSNP01) and incubated on ice for 30 min. For mechanical fragmentation, the nuclei suspension was sonicated in an ice-water bath using a Vibro Cells sonicator (Sonics) equipped with 3 mm microtip at 33% amplitude with 3 s ON / 5 s OFF pulse setting for a total of 1 min.

#### High salt-based nuclear lysis

The nuclei pellet was resuspended in 100 μL of high-salt buffer: 10 mM HEPES pH 7.5, 800 mM NaCl. The suspension was mixed well by pipetting until it became noticeably viscous. 1 unit of SuperNuclease was then added and the sample was further incubated on ice for 30 min until the viscosity had completely disappeared.

To improve the cryoEM behaviour of the sample after high-salt buffer lysis, the salt concentration was then reduced to physiological levels by either salt dilution or chromatin harvesting. In the salt-dilution approach, the sample was diluted fivefold using low-salt buffer (20 mM HEPES pH 7.5, 50 mM NaCl, 0.1% (v/v) Triton X) upon addition of 1 unit of SuperNuclease. The sample was further incubated on ice for 30 min. For chromatin harvesting, the chromatin was collected by centrifugation at 18,800*g* for 10 min at 4 °C. The supernatant was carefully removed and the chromatin was digested with 1 unit SuperNuclease in 100 μL of buffer: 20 mM HEPES pH 7.5, 110 mM KCl, 10 mM NaCl, 0.5 mM MgCl_2_, 1 % (w/v) Trehalose.

### CryoEM sample preparation

All cryoEM work was carried out in the cryoEM facility of the Department of Biochemistry, University of Cambridge.

For every lysis condition, 3.5 μL of lysate sample was applied to glow-discharged Quantifoil R2/2 (300 mesh Cu) grids, either with or without an additional 2 nm continuous carbon support film. Grids were blotted for 5 s at 8 °C, 100% humidity and flash frozen in liquid ethane using Vitrobot Mark IV (ThermoFisher Scientific) equipped with Reverse-Action Tweezer (Nanosoft). The grids were screened in a 200 kV Talos Arctica microscope equipped with Falcon 3EC detector. For each lysis conditions, about 1000 movies were collected for initial data processing at 73,000 magnification (1.37 Å/pix) with defocus values ranging from -3.0 to -1.5 μm.

High-resolution data collection was performed for a sample of nuclear lysate prepared from high-salt lysis and chromatin harvesting protocol, on a 300 kV Titan Krios microscope equipped with Falcon 4i detector and SelectrisX energy filter. Automatic data collection was performed using EPU software version 3.14 (Thermofisher) with defocus values from -2.0 to -0.8 μm. 6305 movies were collected at 135,000 magnification (0.929 Å/pix) with a total electron dose of 47.36 e/Å^2^.

### CryoEM processing

All image processing was performed in cryoSPARC v4.7.1^15^.

#### Talos dataset processing

For all screening datasets collected on the 200 kV Talos microscope, movies were processed using Patch Motion Correction and Patch CTF Estimation and denoised with Micrograph Denoiser. Particles were picked using the Blob Picker on denoised movies, followed by particle extraction. The quality of the dataset was judged from the output of 2D classification (200 classes, 50 online-EM iterations, batch size of 500 per class).

For datasets where 2D classes showed clear evidence of the presence of nucleosome particles, the particle set was improved by iterative 2D classification and then used to generate 3D Ab Initio models, which were refined by heterogeneous refinement. 3D classes that resembled nucleosome core particles were further refined with homogeneous refinement. A detailed description of the Talos dataset processing steps is provided in Supplementary Figure 1 and 2.

#### Krios dataset processing

For the 300 kV Krios dataset, movies were processed using Patch Motion Correction and Patch CTF Estimation and denoised using Micrograph Denoiser. 1,183,042 particles were picked using the Blob Picker on denoised movies, followed by particle extraction with box size 128 pix (3.02 Å/pix) and 2D classification (200 classes, 50 online-EM iterations, batch size of 500 per class). An initial set of 274,321 nucleosome particles selected from 2D class screening was used for a round of template-based picking and also to generate a 3D model using Ab initio Reconstruction.

4,935,164 particles were picked from template-based picking and extracted at box size 128 pix (3.02 Å/pix). The dataset was split into five subsets of around 1 million particles each with Particle Set and subjected to 2D classification. To enhance signal-to-noise, the previous 274,321 particles used for Ab initio Reconstruction were supplemented to each subset during 2D classification. Good 2D classes from all subsets were selected and pooled, and duplicate particles were removed using Remove Duplicates (Minimum separation distance of 50 Å), resulting in a curated set of 786,762 particles (Supplementary Figure 3).

The 786,762 particles were further subjected to heterogeneous refinement using the models from the ab initio reconstruction. The best 3D classes were re-extracted at box size 384 pix (1.006 Å/pix) and subjected to Non-uniform refinement^16^. The map resolution was further improved by applying corrections for per-particle defocus and third-order aberrations (Tilt and Trefoil)^17^. Low-scale particles with Per-particle scale value ≤ 0.99 were removed, and the remaining 286,667 particles were subjected to polishing and re-extracted without binning using Reference-based Motion Correction. A final step of Non-uniform Refinement with optimization for per-particle defocus and higher order aberrations yielded a map at resolution of 2.3 Å (Supplementary Figure 3), which was post-processed with DeepEnhancer^18^. Particle-orientation diagnostics and the local resolution of the cryoEM map are shown in Supplementary Figure 4.

For map inspection purpose, the cryoEM map was cropped in real space to 180 pix (0.929 Å/pix) using the Volume Tools utility. The PDB entry 6IPU for the high-resolution X-ray crystal structure of the human nucleosome core particle^19^ was rigid-body fitted into the post-processed map, and refined using ServalCat^20^. Structural drawings were generated using ChimeraX 1.10^21^.

### Cartoon image generation

Cartoon drawings were prepared with Nano Banana 2 or Nano Banana Pro (Google) and Inkscape v1.4.3 (https://inkscape.org/).

## Results

### Screening nuclear lysis condition for cryoEM imaging

To demonstrate the feasibility of cryoEM single particle analysis on samples of nuclear extracts, we focused on the nucleosome core particle, the fundamental constituent unit of chromatin and an abundant nucleoprotein complex in the nucleus. We first synchronized the HEK293T cells at the G1/S boundary using a double thymidine block to ensure a homogeneous state of the nuclear extracts.

Nuclei were isolated after cell lysis using hypotonic buffer and the cytoplasmic content was removed to prevent cytoplasmic macromolecular assemblies from dominating particle picking and 2D classification during cryoEM processing. We evaluated two distinct nuclear lysis methods, where lysis was achieved either by the use of detergent or high salt (Figure 1).

Lysates were then applied to cryoEM grids with either open holes or a thin continuous carbon coating. Initial screening of grids prepared using open-hole grids showed mostly aggregation of the nuclear extract. However, minimal aggregation was observed with nuclear extracts that were vitrified using carbon-coated grids (Supplementary Figure 5). It is likely that the presence of the carbon support keeps the particles away from the air-water interface by providing a stable absorption surface^22,23^.

Due to the macromolecular crowding of the nuclear extract, it was difficult to assess the quality of the sample based solely on inspection of the micrographs. Therefore, for each lysis methods we collected ∼1000 movies using 200 kV Talos microscope from carbon-coated grids, followed by cryoEM data processing. To improve particle picking in the crowded environment, the micrographs were denoised using cryoSPARC’s Micrograph Denoiser.

Of the two methods, only high-salt lysis yielded 2D classes containing clearly recognisable NCPs (Figure 2A), whereas we couldn’t reliably identify any NCP or other macromolecular particle in the 2D classes of the sample lysed with detergent-based buffer (Supplementary Figure 6). Subsequent 3D reconstruction of the particles from the high-salt lysis condition yielded a low-resolution cryoEM map which confirmed the identification of the NCP in the sample. It is likely that the high-salt concentration in the lysate elevated background scattering in the vitreous ice, resulting in a poor signal-to-noise ratio and compromising accurate particle alignment and resolution of the map.

**Figure 2.**
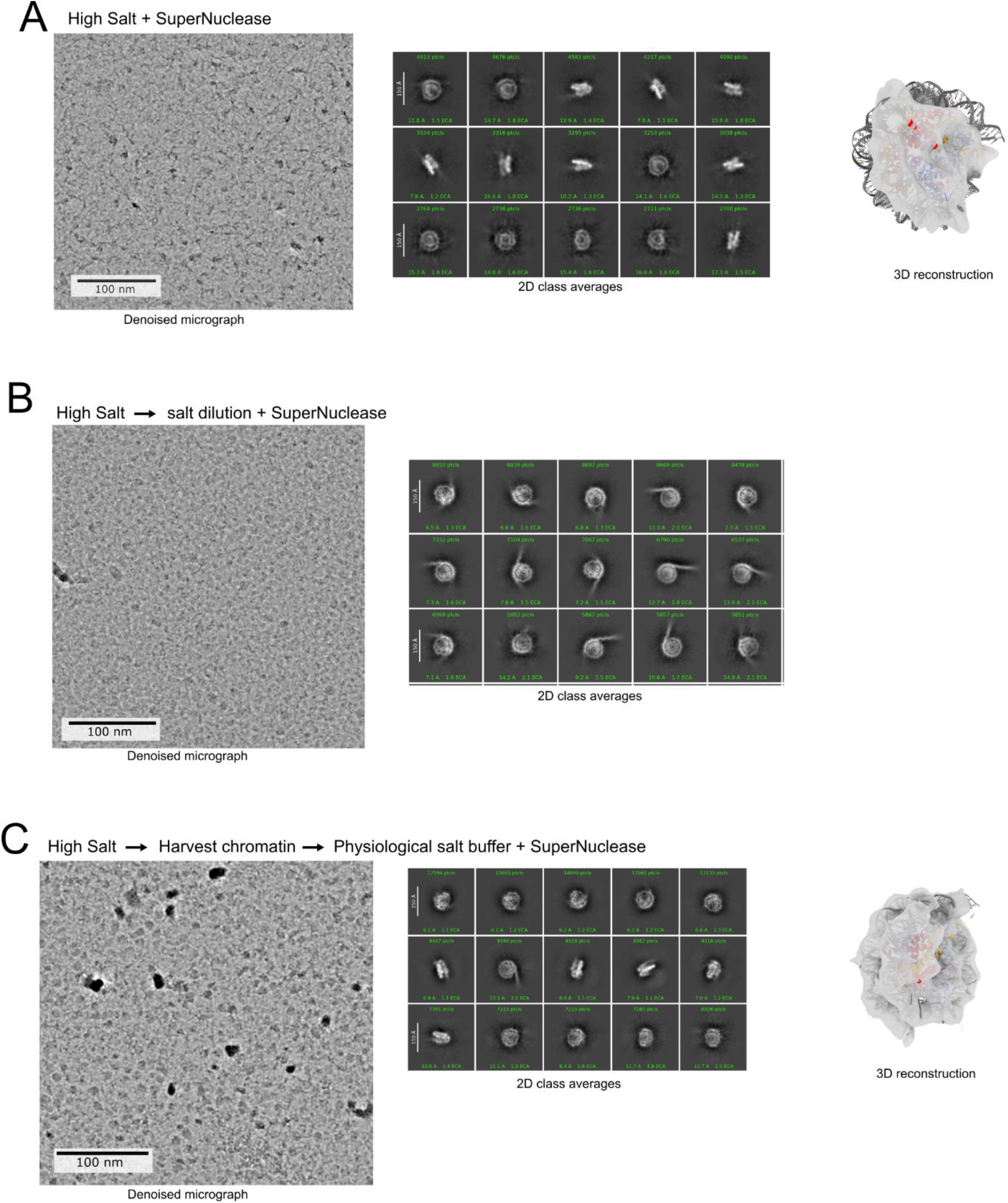
Initial CryoEM screening of the nuclear extract prepared from high-salt lysis approaches. Representative denoised micrographs of carbon-coated Quantifoil grid and 2D class averages for (A) high-salt lysis with SuperNuclease digestion, (B) diluted high salt lysate with SuperNuclease digestion, and (C) harvested chromatin from high-salt lysis with SuperNuclease digestion. 3D reconstruction of the samples shown in (A) and (C), with PDB model 6IPU fitted into the corresponding density maps.

To improve the signal-to-noise ratio, we first reduced the salt concentration in the high-salt lysate by five-fold dilution. This significantly improved the particle alignment and yielded 2D classes showing clear secondary structure features of the NCP. However, the particles exhibited severe preferential orientation biased towards top views, thereby preventing reliable 3D reconstruction (Figure 2B).

Further decisive improvement was obtained through harvesting the chromatin by centrifugation and resuspending it in buffer containing physiological salt concentration and Trehalose as cryoprotectant^24,25^; the extra step alleviated the preferential orientation of the particles and yielded 2D classes with a range of particle orientations (Figure 2C). 3D reconstruction of the NCP from the screening dataset of the chromatin-harvesting sample showed visible major and minor grooves of the DNA in the cryoEM map. Subsequent high-resolution data collection yielded a reconstruction of the nucleosome core particle at 2.3 Å (Figure 3A).

**Figure 3.**
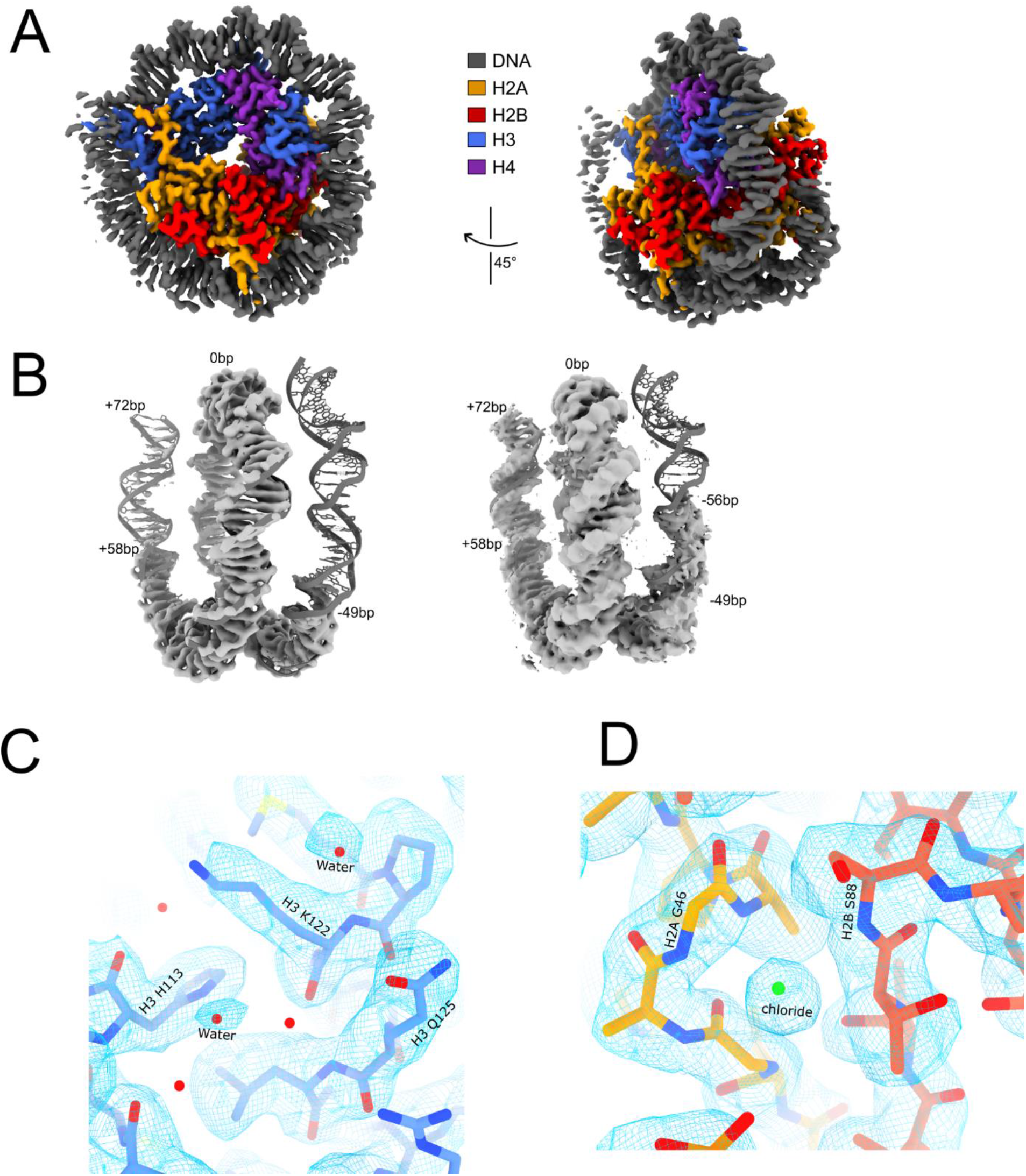
CryoEM density map of the nucleosome. (A) Two rotated views of the cryoEM density map for the nucleosome core particle from nuclear cell extracts. The density corresponding to DNA, H2A, H2B, H3, and H4 is coloured grey, orange, red, blue, and purple, respectively. (B) Portion of the cryoEM density map for the nucleosomal DNA with superimposed ribbon model of the DNA from PDB entry 6IPU. Two different contour levels of the map are shown (higher on the left, lower on the right). (C) Superposition of PDB entry 6IPU with the CryoEM map of the nucleosome core particle obtained from nuclear extracts of HEK293T cells. (D) Detail of the cryoEM showing a density peak that was attributed to a chloride ion (green sphere) in the PBD entry 6IPU.

### Structural features of the NCP from nuclear extracts

The high-resolution crystal structure of the human NCP^19^ (PDB:6IPU) was readily fitted into the cryoEM density map. Comparison of the deposited crystal structure with the model that was real-space refined against the cryoEM map showed remarkably good agreement (a root mean square deviation (RMSD) of ∼0.3 Å for all C_α_ atoms and RMSD of ∼1.0 Å for DNA). The DNA model from the crystal structure fit well to the density; however, only about 128 bp of the DNA was observed as opposed to the canonical ∼147 bp DNA (Figure 3B). The DNA in the NCP structure reported here derives from endogenous genomic DNA, rather than the unique nucleotide sequences – such as the Widom 601 sequence – that are commonly used in reconstituted NCP for structural studies. The genomic DNA may bind to the histone octamer less tightly, thus leading to the observed unwrapping of the ends of nucleosomal DNA. Similar to a previously reported cryoEM study, the DNA unwrapping observed here is predominantly asymmetrical, with one DNA arm being more resolved than the other^26^. However, we cannot exclude that the high-salt extraction protocol further might have contributed to DNA unwrapping, as a high concentration of NaCl is known to induce NCP disassembly^27^.

The high-resolution cryoEM map also allow the visualisation of several ordered water molecules in agreement with the crystal structure (Figure 3C). Interestingly, we also observed an obvious cryoEM density in place, which was modelled as a chloride ion in crystal structures (Figure 3D). While we cannot rule out that some chloride ions were introduced during the high-salt lysis approach, the same chloride ion is observed in high resolution crystallographic models. This chloride ion is stabilised by electrostatic dipole interaction from N-terminal α-helices at the H2A-H2B dimer interface, and the chloride ions can be found in both copies of H2A-H2B dimers of the nucleosome.

## Discussion

The *ex-vivo* lysate approach for high resolution cryoEM of nuclear assemblies presented here provides a practical and accessible alternative to FIB milling. By combining high-salt extraction with carbon-coated grids, we obtained a 2.3 Å structure of the nucleosome core particle directly from the nuclear lysate, bypassing the need for physical sectioning, while achieving a comparable resolution to that of purified native human nucleosome structures^28,29^. The entire extraction and vitrification workflow takes less than two hours and is accessible to any laboratory with standard cryoEM grid preparation equipment. While this procedure does work well to resolve the high-resolution structure of nucleosome, it may not work well for transient and fragile nuclear complex, as the high salt treatment dissociates the nuclear lamina, linker histone, and many chromatin-associated factors.

Attempt to use a different approach using the detergent lysis methods did not yield high-quality 2D class averages (Supplementary Figure 6). The detergent-based lysis method has been used widely in immunoprecipitation and pulldown assays^25,30^. SDS PAGE showed similar overall protein profile across all lysate approaches (Supplementary Figure 5D). It is possible that the gentle detergent-based lysis might produce a biochemically heterogeneous sample that retains a vast array of nuclear protein complexes, and that this heterogeneity generates excessive compositional variability that overwhelms SPA classification and may explain the inability to identify nucleosome 2D classes. Incorporation of template-based particle identification using 2D-template matching (2DTM) might help overcome this challenge by picking desired macromolecular assemblies from the crowded nuclear lysate^31,32^. Alternatively, subtomogram averaging of tilt series data collection of the nuclear lysates could also help overcome the crowding and heterogeneity of the lysate.

Here we have shown that the *ex vivo* lysate approach as promising accessible method that bridge between the structural studies of a fully reconstituted *in vitro* system and *in situ* cryoET. The approach combines near physiological biochemical context with the speed, resolution and accessibility required for broader adoption in structural nuclear biology and beyond. The cryoEM density map determined here achieved sufficiently high resolution to identify water and chloride ions density, demonstrating the potential of this method for small molecule screening directly from nuclear extracts. Given the simplicity of the workflow, this method has the potential to enable rapid structural screening of nucleosomes under a range of conditions, and to capture their interactions with other nuclear factors.

Ultimately, the success of this approach opens the door to the characterisation of other macromolecular assemblies that localise to chromatin. Thus, the challenge for the future is to extend the method’s applicability to the structural analysis of other abundant nuclear complexes with important roles in DNA replication, transcription and repair.

## Acknowledgements

We would like to thank Dimitri Chirgadze and staff at the cryoEM Facility of the Department of Biochemistry for help with data collection. This work was funded by Wellcome Trust award 221892/Z/20/Z to LP.

## Conflict of interest

The authors declare no conflict of interest.

### Author contributions

DSK performed all the experiments, HA maintained the cell culture and performed the nuclear lysis optimization, DSK and LP wrote the paper with input from all authors, LP directed the research.

### Data Availability

The cryoEM map of the NCP has been deposited in the EMDB as entry EMD-58008. The micrographs have been deposited in EMPIAR as entry EMPIAR-13614.

