## Supplementary Figures for "High-resolution cryoEM of nucleosomes in nuclear extracts of mammalian cells"

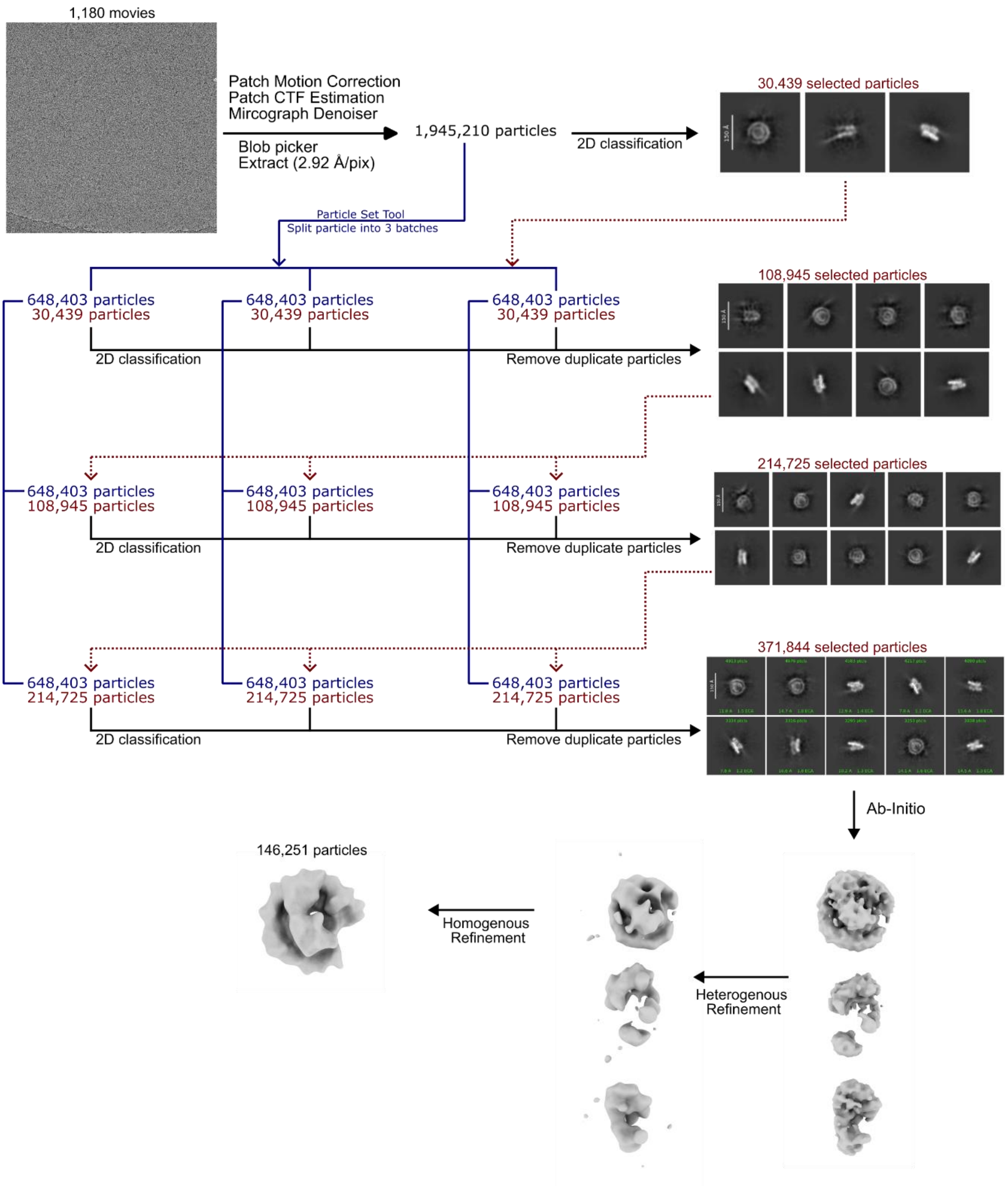


**Supplementary Figure 1: CryoEM data processing workflow for Talos dataset collected from high-salt nuclear lysate.** After motion correction, CTF estimation, micrograph denoising, blob-based particle picking and 2D classification, a subset of 30,439 particles that resembles nucleosome was selected. The 1,945,210 particles that was blob picked was further divided into three subsets, and each underwent 2D classification together with 30,439 particles that resembles nucleosome particles. Nucleosome-like 2D classes from all subsets were selected and pooled, and duplicate particles were removed. This process was done iteratively for three times, which yielded 371,844 final particles. This final particles was used for ab initio reconstruction and heterogeneous refinement. The best 3D class was then subjected to homogenous refinement to yield the final map.

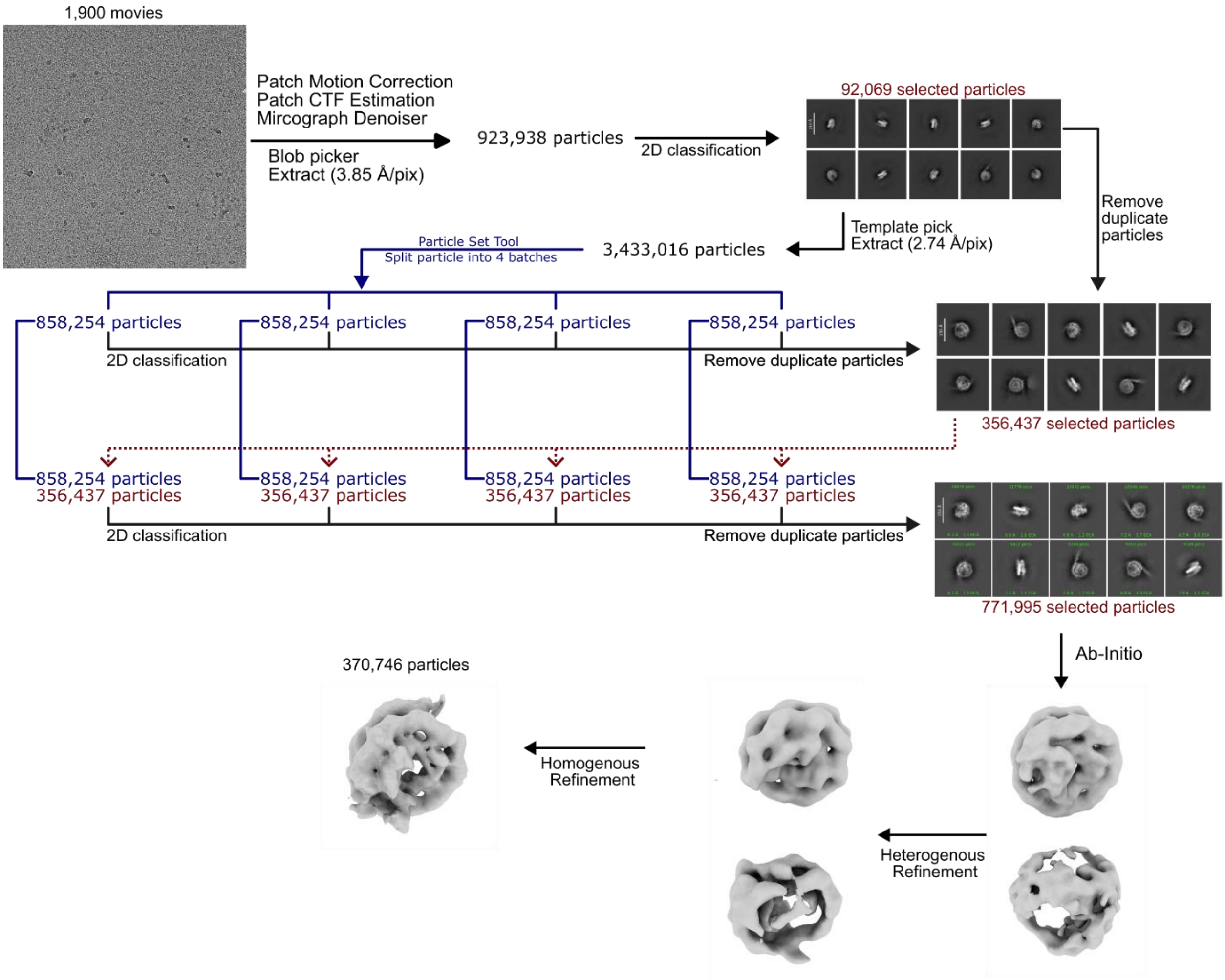


**Supplementary Figure 2: CryoEM data processing workflow for Talos dataset collected from optimized nuclear extraction.** After motion correction, CTF estimation, micrograph denoising, blob-based particle picking and 2D classification, this resulted in a subset of 92,069 particles that resembles nucleosome. 3,433,016 particles were picked via template-based picking and they were further divided into four subset and were subjected to 2D classification independently. All nucleosome-like 2D classes were selected and pooled together with the initial 92,069 particles, and duplicated particles were removed. One more round of 2D classification was done by supplementing the 356,437 nucleosome-like particles into each 2D classification set, to isolate more nucleosome-like particles from the dataset. The final 771,995 particles were used for ab initio reconstruction and heterogeneous refinement. The best 3D class was then subjected to homogenous refinement to yield the final map.

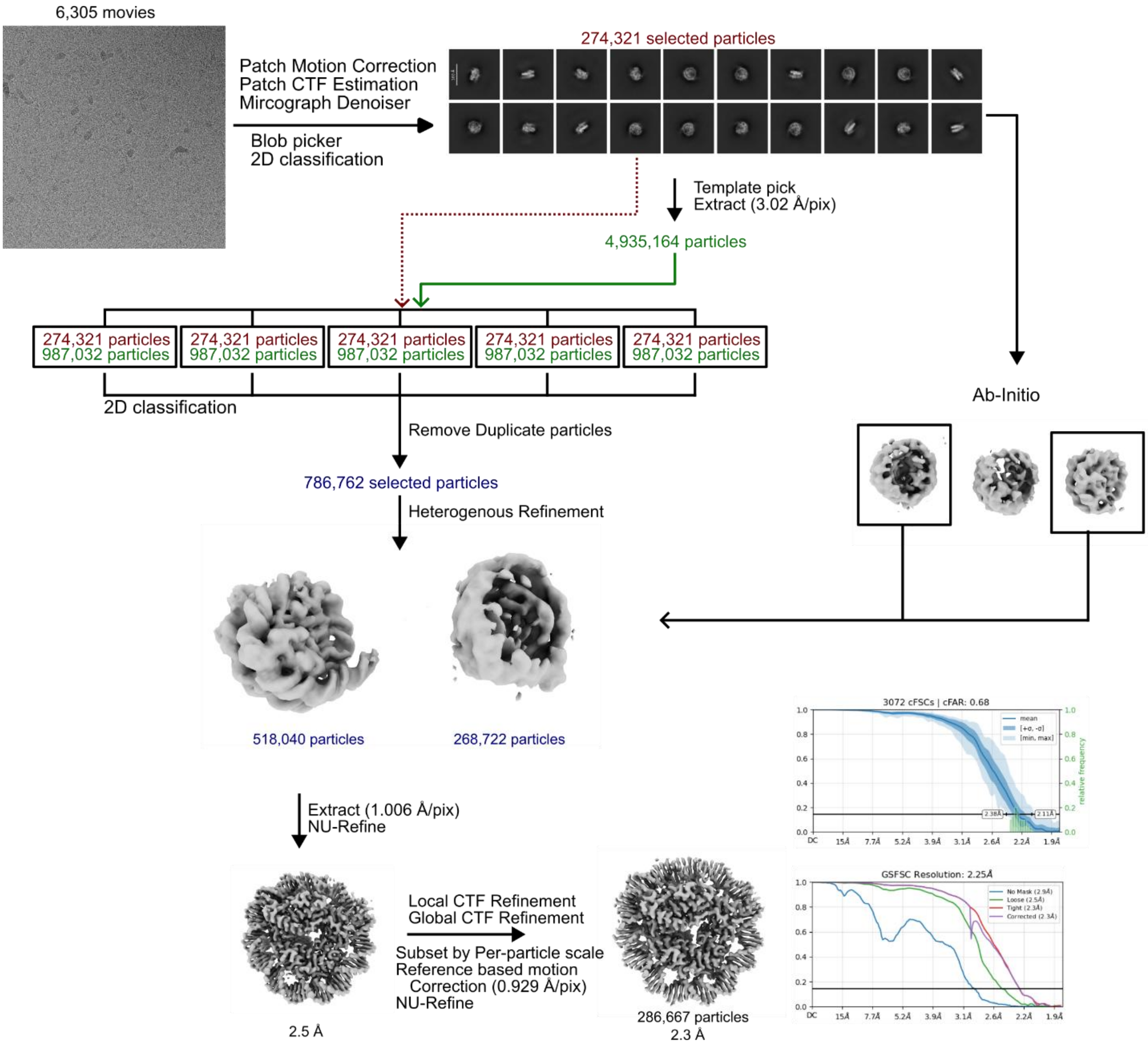


**Supplementary Figure 3: CryoEM data processing workflow for Krios dataset collected from optimized nuclear extraction.** After motion correction, CTF estimation, micrograph denoising, blob-based particle picking and 2D classification, this resulted in a subset of 274,321 particles that resembles nucleosome. 4,935,164 particles were picked via template-based picking and they were further divided into five subsets. Each of the subset were further supplemented with 274,321 particles and underwent 2D classification independently. All nucleosome-like 2D classes were selected and pooled together, and duplicated particles were removed. The final 786,762 particles were used for heterogeneous refinement using the 3D initial model generated from the initial 274,321 good particles. The best 3D class was then subjected to non-uniform refinement to yield a high resolution nucleosome map. The resolution of the map was further enhanced by performing local CTF refinement, global CTF refinement and reference-based motion correction.

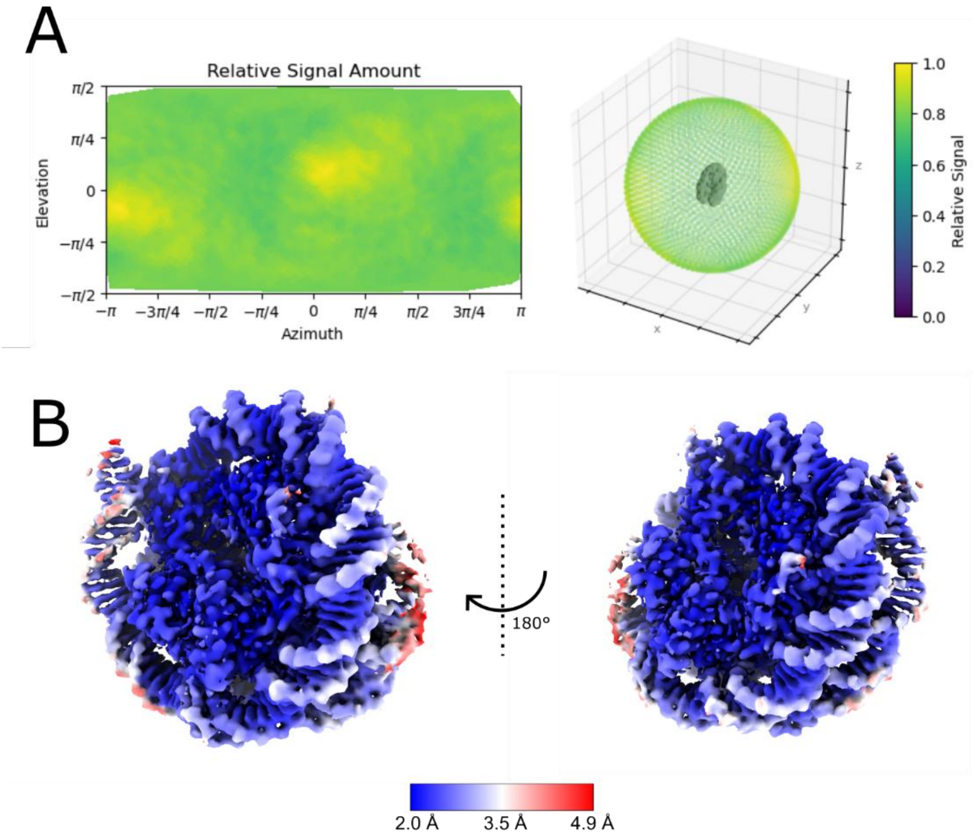


**Supplementary Figure 4: Directional distribution and local resolution of the cryoEM map.** (A) Plot of relative signal versus viewing direction, together with a 3D scatter plot of particle orientations. (B) Local resolution map displayed from with two different views.

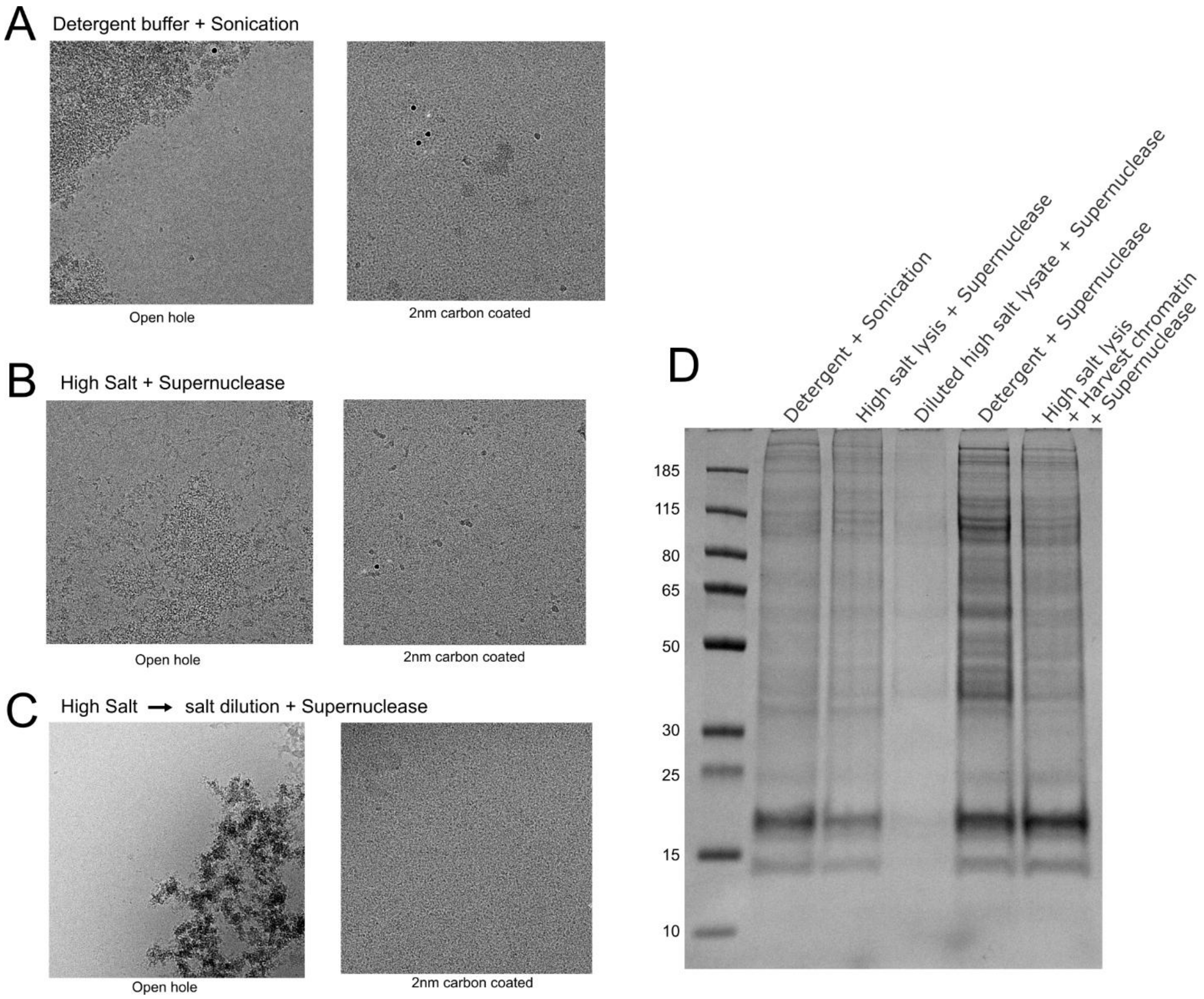


**Supplementary Figure 5: CryoEM Talos screening of different grid types.** Representative micrograph of open hole and 2nm carbon-coated Quantifoil grids for (A) sonication, (B) high salt lysis with SuperNuclease digestion, and (C) diluted high salt lysate with SuperNuclease digestion. (D) SDS PAGE of the cryoEM samples prepared using different lysis approaches.

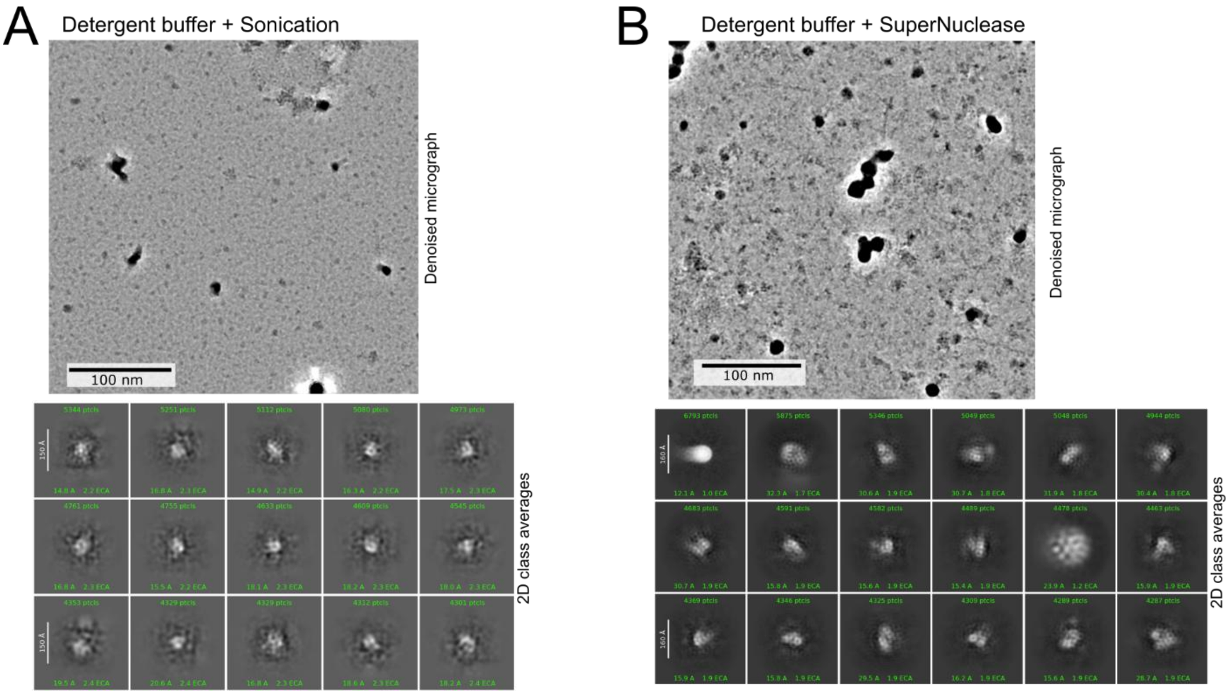


**Supplementary Figure 6: CryoEM Talos screening of the nuclear extract from detergent lysis approaches.** Representative denoised micrographs of carbon coated Quantifoil grid and 2D class averages for (A) detergent lysis with sonication and (B) detergent lysis with SuperNuclease digestion.

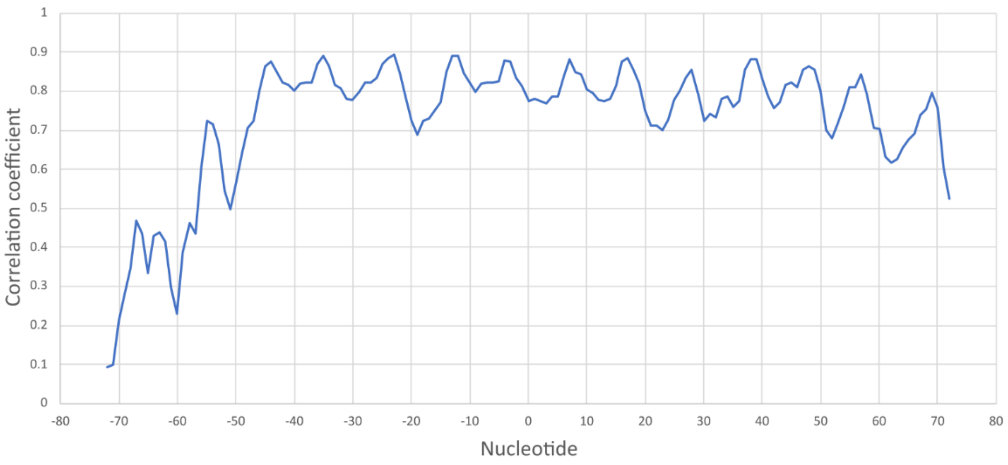


**Supplementary Figure 7: Map-to-model correlation plot of the nucleotide chain in cryoEM map.**
